# Synthesis and Characterisation of a Macrophage-derived Hybrid Nanoparticles for Doxorubicin Delivery to Glioblastoma

**DOI:** 10.64898/2026.05.20.726551

**Authors:** Girstautė Dabkevičiūtė, Christian Celia, Vilma Petrikaitė

## Abstract

Glioblastoma (GBM) presents significant therapeutic challenges due to its aggressive nature, complex microenvironment and the limitations of conventional drug delivery systems. In this study, hybrid nanoparticles were developed by combining synthetic liposomes with macrophage-derived extracellular vesicles (EVs) to harness the strengths of both platforms. Two distinct liposomal formulations, DPPC:Chol:DSPE-mPEG2000 (F1) and DPPC:DPPS:Chol:DSPE-mPEG2000 (F2), were used as the basis for the synthesis. EVs derived from J774 macrophages were integrated with F1 and F2 to create hybrid nanoparticles (H-F1 and H-F2). Doxorubicin (DOX) was encapsulated using a pH gradient and a remote loading procedure. The mean particle size of H-F1-DOX and H-F2-DOX was 158.2 ± 1 nm and 162.8 ± 9 nm, respectively. The polydispersity index (PDI) was 0.130 ± 0.012 and 0.084 ± 0.033, while the zeta potential values were −14.9 ± 0.7 mV and −26.7 ± 3.1 mV, respectively. H-F2-DOX exhibited the highest encapsulation efficiency (EE%), reaching 76.5±3.4%. The encapsulated hybrids remained stable up to one week, at +5°C. The release of DOX from H-F2-DOX in DMEM supplemented with 10% serum showed pH sensitivity, with total DOX release of 64.9 ± 5.3% at pH 7.4 and 90.7 ± 6.5% at pH 5.5. The cell viability assay demonstrated that all formulations exhibited strong cytotoxic effects against GBM cells under normoxic conditions, with H-F2-DOX showing the most potent effect under hypoxia-mimetic conditions.

## 1. Introduction

Glioblastoma multiforme (GBM) is the most lethal brain cancer in adults, accounting for approximately half of all primary malignant brain tumors [1]. It remains incurable, mainly due to biochemical and biophysical barriers to efficient drug delivery to the tumor site [2]. In particular, the blood-brain barrier (BBB) inhibits transport of more than 98% of small-molecule drugs and all macromolecule drugs into the brain [3]. Alongside the BBB, the extracellular matrix (ECM), rich in hyaluronic acid, hinders further penetration of therapeutics, contributing to a protective and supportive environment for GBM [4]. Thus, overcoming ineffective pharmacotherapy requires the development of safe and novel strategies to enhance drug concentrations within the tumor, which are limited by the complex GBM microenvironment, including its protective barriers.

Nanoparticles have shown great potential for cancer treatment due to their ability to improve drug pharmacokinetics, reduce systemic toxicity, and increase tumor targeting [5]. The success of nanoparticle-based therapeutics is demonstrated by the clinical approval of several nanoformulations, primarily liposomes, for the treatment of Kaposi’s sarcoma, ovarian cancer, metastatic breast cancer, and other malignancies [6,7,8,9]. However, most clinical trials investigating nanoparticles have failed to yield substantial benefits for GBM patients, highlighting the persistent challenges of GBM treatment [10]. Known limitations of nanoparticle delivery in GBM include heterogeneous passive diffusion through the BBB, reduced tissue penetration due to the ECM and premature uptake by tumor-associated macrophages (TAMs) [11,12,13]. Thus, passive targeting of nanoparticles remains limited and inconsistent, and active targeting is being explored as a more promising approach [11,14]. In GBM, active targeting typically involves surface modifications with ligands that guide nanoparticles to specific receptors in the BBB and/or malignant cells, such as peptides, antibodies, aptamers, sugars, and small molecules like folic acid [15,16,17,18]. However, for effective targeted delivery, single ligand conjugation might be insufficient, while the chemistry of multiple ligand modifications is technically challenging [19]. Therefore, extracellular vesicles (EVs), as natural cargo carriers with complex membrane architectures that can facilitate targeted delivery, have gained increasing attention as promising drug delivery systems [20,21].

EVs represent a heterogeneous population of naturally occurring vesicles that facilitate communication between cells [22,23,24]. Being released by almost all cell types, EVs differ in size and biogenesis. This includes small endosome-derived particles (exosomes) and larger vesicles generated by budding from the plasma membrane [25,26]. Notably, these vesicles can cross the BBB, making them promising candidates for the treatment of neurodegenerative diseases [27,28,29]. Research, specifically on small EVs, has shown interaction with the BBB via transferrin receptors, eliminating the need for additional surface modifications [30]. Several studies have utilized EVs isolated from different commercial GBM cell lines, such as U-87MG, GL261, and U-251MG, due to their inherent “homing” effect toward the tumor [31,32,33]. However, tumor-derived EVs have been reported to reprogram healthy cells, thereby promoting tumor progression through the induction of angiogenesis, immunosuppression, and fibroblast activation [34,35,36,37]. Consequently, immune cells have been recognized as a promising source of EVs for the development of biomimetic delivery systems [38,39].

In this study, we derived EVs from non-polarized macrophages, given the extensive infiltration of immune cells in GBM and their resemblance to the M0 phenotype [40]. Macrophage-derived EVs were fused with two liposomal formulations for doxorubicin (DOX) delivery to GBM cells. Previous studies have shown that encapsulating cargo into EVs alone is challenging without compromising their structural integrity, often leading to low efficiency [41,42,43,44]. Therefore, hybridization was conducted to create stable biomimetic nanoparticles with enhanced cargo encapsulation efficiency and preserved natural targeting properties of EVs. Liposomes used for hybridization were evaluated as bases for hybrid nanoparticles to assess the influence of lipid composition on hybridization, stability, encapsulation efficiency, drug release, and U-87MG cell viability. DOX was selected as the cargo due to its potent *in vitro* anticancer activity against GBM cell lines, limited BBB permeability, and inherent fluorescence enabling easy tracking [45,46].

## 2. Materials and Methods

### Materials

1,2-dipalmitoyl-sn-glycero-3-phosphocholine (DPPC), 1,2-dipalmitoyl-sn-glycero-3-phospho-L-serine (DPPS), 1,2-distearoyl-sn-glycero-3-phosphoethanolamine-N [methoxy (polyethylene glycol)-2000] (DSPEmPEG2000) were purchased from Avanti Polar Lipids Inc. (Birmingham, AL, USA). The cell lines J774 and U-87MG were obtained from American Type Culture Collection (ATCC, USA). DOX hydrochloride was purchased from Abcam (UK).

### Liposome

Liposomes were prepared using the thin layer evaporation (TLE) method with the final lipid molar ratio 6:3:1 for DPPC:Chol:DSPE-mPEG2000 (F1), respectively, and 7:4:6:1 for DPPC:DPPS:Chol:DSPE-mPEG2000 (F2), respectively. Lipids were dissolved in an organic solvent solution (chloroform/methanol, 3:1 v/v) and evaporated using a rotary evaporator IKA RV 10 at 56°C. The remaining solvent was left to evaporate overnight. Then, the resulting lipid film was hydrated with 10 mM HEPES buffer (pH 7.4) for empty liposomes or 250 mM ammonium sulfate (pH 5.5) for therapeutic liposomes to obtain a final lipid concentration of 20 mg/mL. After hydration, the suspension underwent three cycles of warming at 56°C and vortexing at 600 rpm and was left for stabilization at 60°C for 1 hour. Finally, liposomes were extruded through polycarbonate membrane filters ranging from 800 nm to 100 nm, using The Avanti Mini-Extruder (Avanti Polar Lipids Inc., Birmingham, AL, USA) at 60°C.

### GBM cell culturing

GBM cells were cultured in DMEM Glutamax cell culture medium (Gibco, Carlsbad, CA, USA) supplemented with 10% fetal bovine serum (FBS, Gibco) and 1% of 10,000 U/mL penicillin and 10 mg/mL streptomycin solution at 37°C with 5% CO_2_ in a humidified atmosphere. Cells were grown to 70% confluency, trypsinized with 0.125% TrypLE Express (Gibco) before passage and used until passage 20.

### EVs isolation and purification

The EVs were isolated and purified as described previously [47]. Briefly, J774 murine macrophage cells were cultured with DMEM Glutamax cell medium (Gibco, Carlsbad, CA, USA) supplemented with 10% FBS (Gibco) and 1% of 10,000 U/mL penicillin and 10 mg/mL streptomycin solution. After 48 hours of incubation, the cell culture medium was replaced with a fresh one, supplemented with 10% v/v exosome-depleted FBS (Gibco) and incubated for 48 hours. The medium was then collected and centrifuged twice: first at 700 ×g for 5 min and then at 2000 ×g for 10 min, followed by filtration. At the purification step, the medium was concentrated with Centricon^**®**^ Plus-70 centrifugal filters (MiliporeSigma) up to a final volume of 1 mL and loaded into qEV original 70 nm columns (Izon Science). Multiple fractions were collected from the column and characterized in terms of size and visualized by scanning electron microscopy (SEM). Transmission electron microscopy (TEM) characterisation has been reported previously [47], therefore, it is not included in this study. The selected EVs fractions for hybridization were further characterized by dynamic light scattering (DLS).

### SEM

The isolated EVs were visualized using a Hitachi S-3400N scanning electron microscope (Tokyo, Japan). 10 μL of EVs were fixed by adding an equal volume of 4% formaldehyde and incubated for 20 minutes. The samples were then washed three times with PBS and allowed to air-dry. To avoid altering the natural surface structure of the samples, imaging was performed without conductive coating, thereby eliminating potential distortions caused by sputter-coating. A 5 keV accelerating voltage was selected during imaging to improve sensitivity to surface features and reduce charging effects from the electron beam.

### Hybrid nanoparticle synthesis

F1 and F2 lipid films were prepared in the same manner as described in the liposome synthesis section. The lipid films were hydrated with 0.5 mL of EVs diluted with HEPES buffer to a final volume of 2 mL for empty hybrids or diluted with ammonium sulfate for therapeutic hybrids. The hydrated films underwent five freeze–thaw cycles, consisting of freezing at −80°C and thawing at 37°C. Finally, hybrid nanoparticles were extruded through polycarbonate membrane filters ranging from 800 nm to 100 nm, using The Avanti Mini-Extruder (Avanti Polar Lipids Inc., Birmingham, AL, USA) at 37°C.

### Physicochemical characterization

The physicochemical properties, such as particle size, PDI, and zeta potential were determined by DLS using Zetasizer Ultra (Malvern Instruments Ltd, Malvern, UK), using a backscattering detection angle of 173°. 20 μL of nanoparticle suspension was diluted to a final volume of 1 mL, and measurements were performed at 25°C. Zeta potential was calculated from electrophoretic mobility according to the Helmholtz–Smoluchowski equation.

### Storage stability

Storage stability of empty and loaded nanoparticles was evaluated at 5°C in HEPES. Samples for DLS were taken once a week for a month and measured for particle size, PDI, and zeta potential.

### Short-term stability

Short-term stability of empty nanoparticles was evaluated under stressful conditions. 200 μL of nanoparticle suspension was added to 1 mL of cell culture medium supplemented with 10% FBS and incubated for 72 hours at 37 °C under magnetic stirring at 300 rpm. The short-term stability was measured at pH values of 5.5 and 7.4. Samples of 20 μL were taken at 0, 1, 2, 4, 6, 8, 24, 48, and 72-hour time points and were measured for particle size and PDI.

### DOX encapsulation

DOX encapsulation was performed using a pH gradient and a remote loading procedure. After lipid hydration with ammonium sulfate and extrusion, the resulting nanoparticles were centrifuged using Vivaspin® Turbo 15 PES centrifugal concentrators (Sartorius, Germany) for 5 cycles of 10 min at 2950 ×g. 500 μL of nanoparticles in HEPES were collected from the concentrator filters and added to 500 μL of DOX solution (2 mg/mL), then incubated at 60°C under magnetic stirring for 1 hour. For hybrid nanoparticles, the incubation temperature was reduced to 37°C to preserve EVs components. Unentrapped DOX was removed using a Tube-O-DIALYZER ® mini dialysis system (G-Biosciences, USA) equipped with a 15 kDa molecular weight cut-off membrane, under continuous stirring for 12 hours at room temperature. HEPES buffer (10 mM, pH 7.4) was used as the receptor medium. After dialysis, encapsulated nanoparticles were characterized in terms of particle size, PDI, and zeta potential.

DOX encapsulation efficiency (EE%) was determined by fluorescence spectroscopy. 50 μL of nanoparticle suspension was centrifuged for 5 cycles at 16873 ×g for 5 min at 4°C. After centrifugation, the supernatant was carefully removed. The formed pellet was dissolved in 1 mL of methanol. 200 μL of methanol solution was added to microplates for fluorescence measurement using a Fluoroskan Ascent microplate reader (Thermo Scientific, USA). Fluorescence measurements were performed at excitation and emission wavelengths of 485 and 538 nm, respectively.

### DOX release

DOX release was evaluated by dialysis using a Tube-O-DIALYZER^®^ mini dialysis system (G-Biosciences, USA) in cell culture medium supplemented with 10% FBS at pH 5.5 and 7.4, maintained at 37°C under continuous stirring. A volume of 220 μL of nanoparticles was loaded into the dialysis system and immersed in 220 mL of DMEM with 10% FBS to ensure sink conditions. Each time 200 μL of the acceptor medium containing released DOX was withdrawn, it was replaced with 200 μL of fresh acceptor medium. The study was conducted over 72 h, with samples collected at 15 and 30 min and at 1, 2, 4, 6, 24, 48, and 72 h.

### Cell viability assay

GBM cell viability was evaluated using 3-(4,5-dimethylthiazol-2-yl)-2,5-diphenyltetrazolium bromide (MTT, Sigma-Aldrich Co., ≥ 97%) reduction assay. U-87MG cells were seeded in 96-well plates in a volume of 100 μL (4×10^3^ cells/well for normoxic conditions and 8×10^3^ cells/well for hypoxia-mimetic conditions). After 24 hours, cells were treated with different concentrations of DOX-loaded nanoparticles and free DOX solution, corresponding to from 0.0032 μg/mL to 1 μg/mL DOX. For hypoxia-mimetic conditions, nanoparticle and free DOX solutions were prepared with CoCl2 at a concentration of 200 μM. Wells with only medium without cells were used as a positive control, and wells with untreated cells served as a negative control. After 72 hours, cells were incubated with 0.5 mg/mL of MTT solution for 3 hours. The formed formazan crystals were dissolved in 2-propanol. Half maximal effective concentration (EC_50_) values were calculated using the Hill equation.

### Statistical analysis

The data was processed using *Microsoft Excel (Microsoft 365, Version 2603)* and *IBM SPSS Statistics (Version 20)*. Normality of residuals from two-way and one-way ANOVA was evaluated using Q–Q plots. Two-way ANOVA followed by Bonferroni-corrected unpaired two-tailed t-tests was used for comparisons of physicochemical characteristics. One-way ANOVA with Tukey’s post hoc test was applied for the analysis of EE%, EC_50_ and cell viability. Differences were considered statistically significant at p < 0.05.

## 3. Results

### Characterization

Two liposomal formulations (F1 and F2), differing in the molar ratios of DPPC, DPPS, cholesterol, and DSPE-mPEG2000, were prepared and characterized in terms of nanoparticle size, size distribution (PDI), and zeta potential; two-way ANOVA results are presented in **Table 1**, while the compositions of the liposomal formulations are shown in **Figure 1A** and characterization data of liposomes and their hybrid nanoparticles are shown in **Figure 1B**. Following extrusion through polycarbonate membranes, both liposomal formulations resulted in a similar size and PDI < 0.2 (**Figure 1B**). The average size of F1 and F2 was 148.2±6.6 nm and 147.9±4.5 nm, respectively. DLS measurements showed that the average zeta potential of F1 was −7.9 ± 1.9 mV, while that of F2 was −23.3 ± 1.7 mV. To obtain therapeutic liposomes, DOX was encapsulated into liposomes using the pH gradient method, which was done after extrusion. Similarly to liposomes, hybrid nanoparticles (H-F1 and H-F2) were synthesized and then characterized. To generate hybrid carriers, F1 and F2 liposomal lipid films were hydrated with EVs that had been collected and selected based on size. The average size of H-F1 and H-F2 was 145.3±3.6 nm and 139.5±6.5 nm, respectively. The hybrids displayed a slightly lower zeta potential, with a maximum decrease of 5 mV observed for H-F1. The fusion did not hinder extrusion, and both hybrids resulted in a narrow size distribution (PDI < 0.1).

**Table 1.** Significance of the effects of formulation and DOX encapsulation on nanoparticle physicochemical properties.

| Synthesis factor | Size |  | Zeta potential |  |
| --- | --- | --- | --- | --- |
|  | F (df) | P value | F (df) | P value |
| Formulation | 2.158 (3) | 0.133 | 31.692 (3) | <0.001 |
| DOX | 48.776 (1) | <0.001 | 0.005 (1) | 0.943 |
| Formulation*DOX | 3.300 (3) | 0.047 | 3.309 (3) | 0.047 |

F, F-statistic; df, degrees of freedom.

**Fig. 1.**
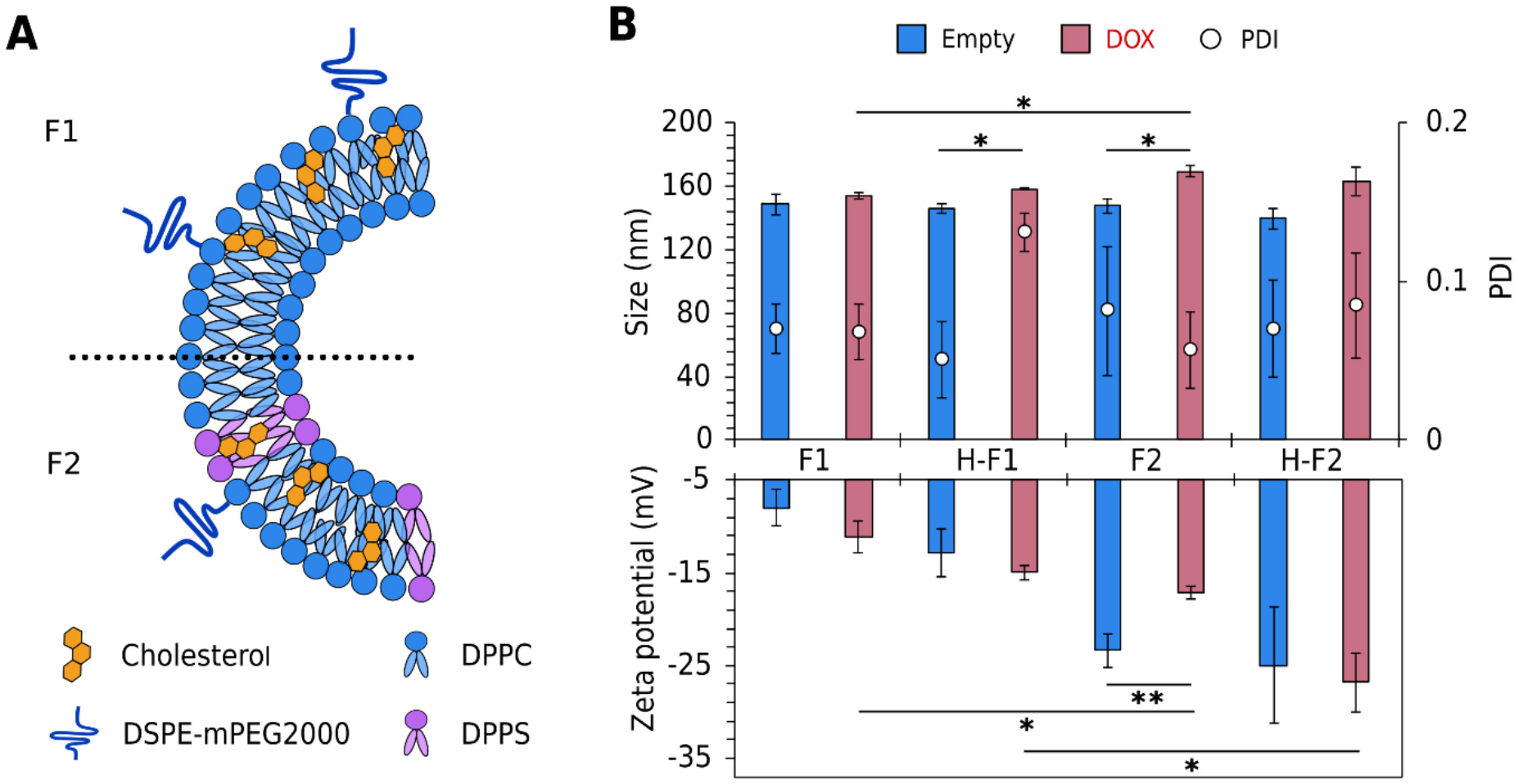
Composition of F1 and F2 liposomes (A) and physicochemical properties of synthesized empty (blue column) and loaded (red column) liposomes and their hybrid nanoparticles (B). Significance levels: p < 0.05 (*), p < 0.01 (**). Statistical analysis was performed using two-way ANOVA followed by post hoc pairwise comparisons using unpaired t-tests with Bonferroni correction. Comparisons between empty formulations are not shown. Data were expressed as mean ± standard deviation (SD), n=3. Abbreviations: DOX, doxorubicin; PDI, polydispersity index.

Two-way ANOVA showed that adding DOX had a significant effect on nanoparticle size, and the effect was dependent on the formulation (**Table 1**). F1-DOX showed minimal change in size, while F2-DOX, H-F1-DOX and H-F2-DOX increased by up to 23 nm compared to the empty nanoparticles. The increase was significant for F2-DOX and H-F1-DOX, but not for H-F2-DOX, after Bonferroni correction (**Figure 1B**). The zeta potential was significantly influenced by the formulation itself, and the interaction between formulation and DOX was also statistically significant (**Table 1**). After encapsulation, F2-DOX exhibited a 6 mV shift in zeta potential, in addition to the observed increase in size. These changes indicate that DOX encapsulation had a greater impact on the properties of F2 than on those of F1 (**Figure 1B**). In comparison among therapeutic liposomes, F2-DOX was significantly bigger and more negative than F1-DOX. When comparing the therapeutic hybrids, their sizes were similar, however, H-F2-DOX exhibited a significantly more negative zeta potential. Finally, after the encapsulation procedure, the PDI of the liposomes and the hybrid nanoparticles were below 0.2.

### Long-term stability

To evaluate the potential shelf-life and ensure reproducibility and data validity, liposomes were stored at +5°C and monitored over one month for physicochemical stability. Both empty and DOX-loaded liposomes were characterized in terms of particle size, homogeneity (PDI), and zeta potential. According to DLS measurements, the average size of empty F1 liposomes increased by approximately 7 nm from week 0 to week 4 (**Figure 2A**). This change was accompanied by a slight increase in PDI, from 0.07 to 0.10. Despite these changes, PDI remained below 0.2, indicating that empty F1 liposomes maintained a narrow size distribution and colloidal stability. Similar to the empty liposomes, the DOX-loaded F1 formulation showed only a minor increase in size over 4 weeks, from 153.7 ± 2.3 nm to 157.1 ± 1.2 nm. The PDI of DOX-loaded F1 liposomes remained below 0.2. After 4 weeks, the zeta potential of empty and DOX-loaded F1 liposomes showed minimal changes compared to week 0, with differences of only 0.4 mV and 2.2 mV, respectively (**Figure 2A**).

**Fig. 2.**
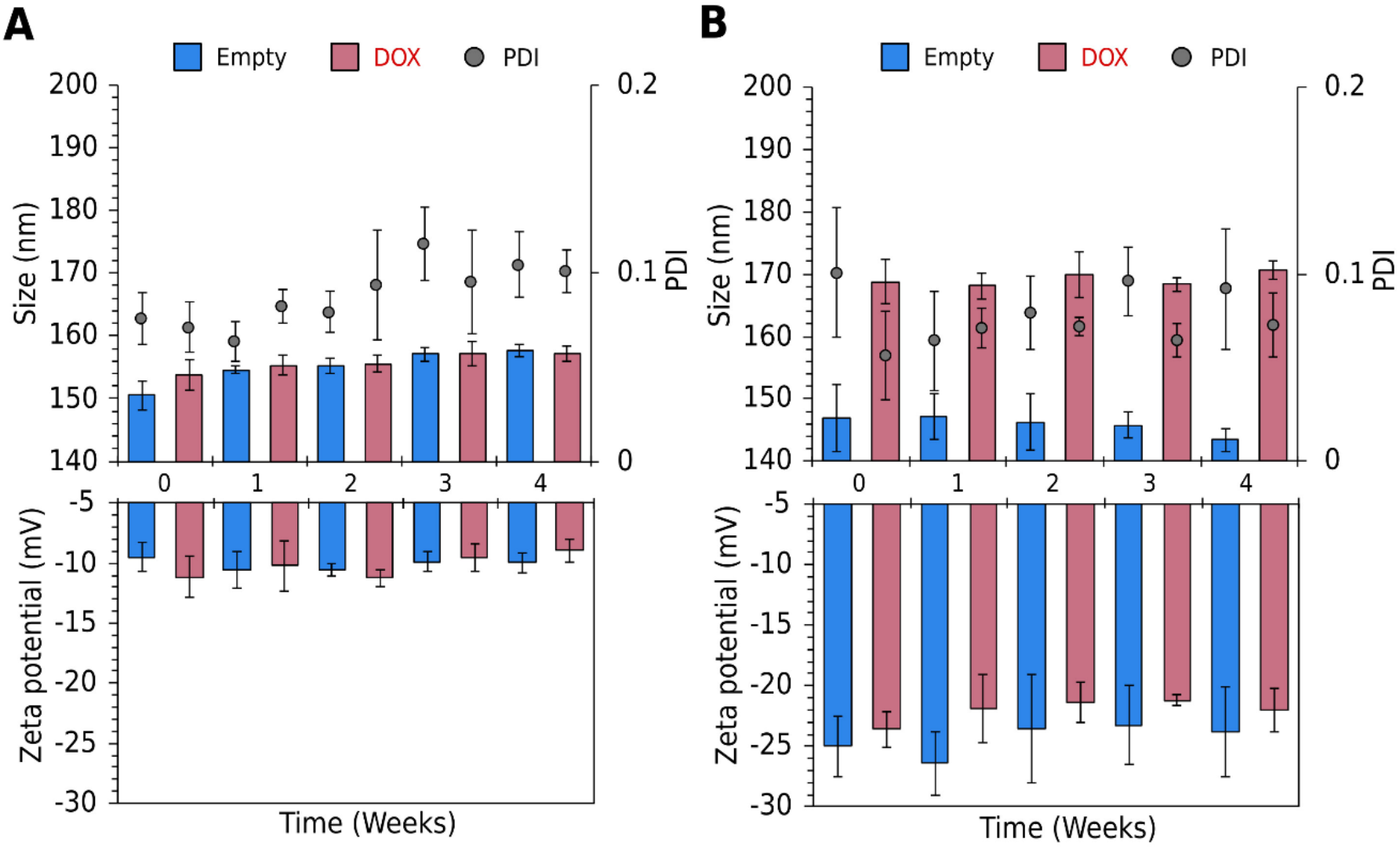
Long-term stability of empty and loaded F1 formulation (A) and F2 formulation (B). Data were expressed as mean ± standard deviation (SD), n=3. Abbreviations: DOX, doxorubicin; PDI, polydispersity index.

Similarly, empty F2 liposomes also remained stable during the storage period, with only a slight change in average size of 2.3 nm between week 0 and week 4 (**Figure 2B**). The PDI remained below 0.2 throughout the storage period, suggesting that the empty F2 liposomes, similarly to F1, maintained a narrow size distribution. DOX loading did not affect the 1-month stability of F2 liposomes, as size changes remained minimal, with an increase of only up to 2 nm over the storage period. PDI for DOX-loaded F2 varied slightly but stayed below 0.1, likewise, the zeta potential remained stable throughout storage.

Physicochemical stability of hybrid nanoparticles was evaluated under the same conditions as for the parent liposomes to determine whether hybridization impacts storage stability. Results have shown that empty hybrid nanoparticles retained their size for a whole month, similarly to the empty liposomes (**Figure 3**). The size distribution remained narrow with a PDI below 0.2. Consistent with stable size and lack of aggregation, zeta potential changes were ≤ 4 mV. However, both hybrid formulations showed a clear shift in size and PDI at week 2 following DOX encapsulation. The hydrodynamic diameter reached ~190 nm, and the PDI exceeded the 0.2 value, suggesting particle aggregation. At week 2, the zeta potential decreased by 7 mV for H-F1-DOX (**Figure 3A**) but increased by 2 mV for H-F2-DOX (**Figure 3B**). Overall, the long-term stability of empty hybrid nanoparticles was comparable to that of their parent liposomes. However, DOX encapsulation strongly influenced the size and size distribution of the hybrid nanoparticles, and aggregation became apparent within one week of storage.

**Fig. 3.**
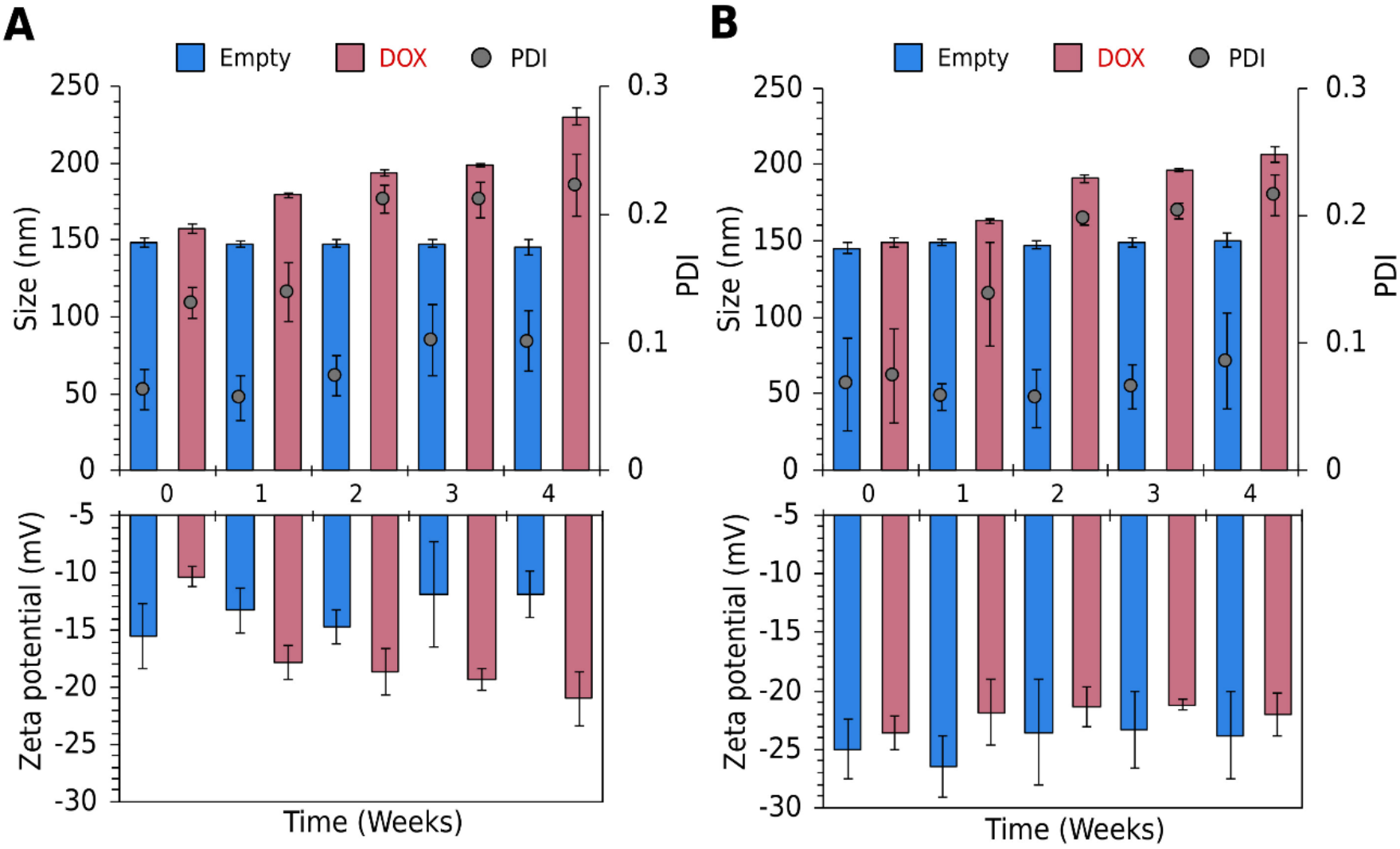
Long-term stability of empty and loaded H-F1 formulation (A) and H-F2 formulation (B). Data were expressed as mean ± standard deviation (SD), n=3. Abbreviations: DOX, doxorubicin; PDI, polydispersity index.

### Short-term stability

Short-term stability of empty nanoparticles was assessed in DMEM containing 10% FBS at neutral and acidic pH for 72 h at 37°C. Samples were continuously stirred throughout incubation to impose stress. After incubation in cell culture medium, both formulations showed a decrease in average size of up to 26 nm, while PDI remained unchanged (**Figure 4A, B**). Despite this reduction, no aggregation or structural destabilization was observed over 72 hours at pH 7.4. At pH 5.5, the size increased for both formulations, peaking at 48 hours, reaching 128.8 nm for F1 and 137.0 nm for F2.

**Fig. 4.**
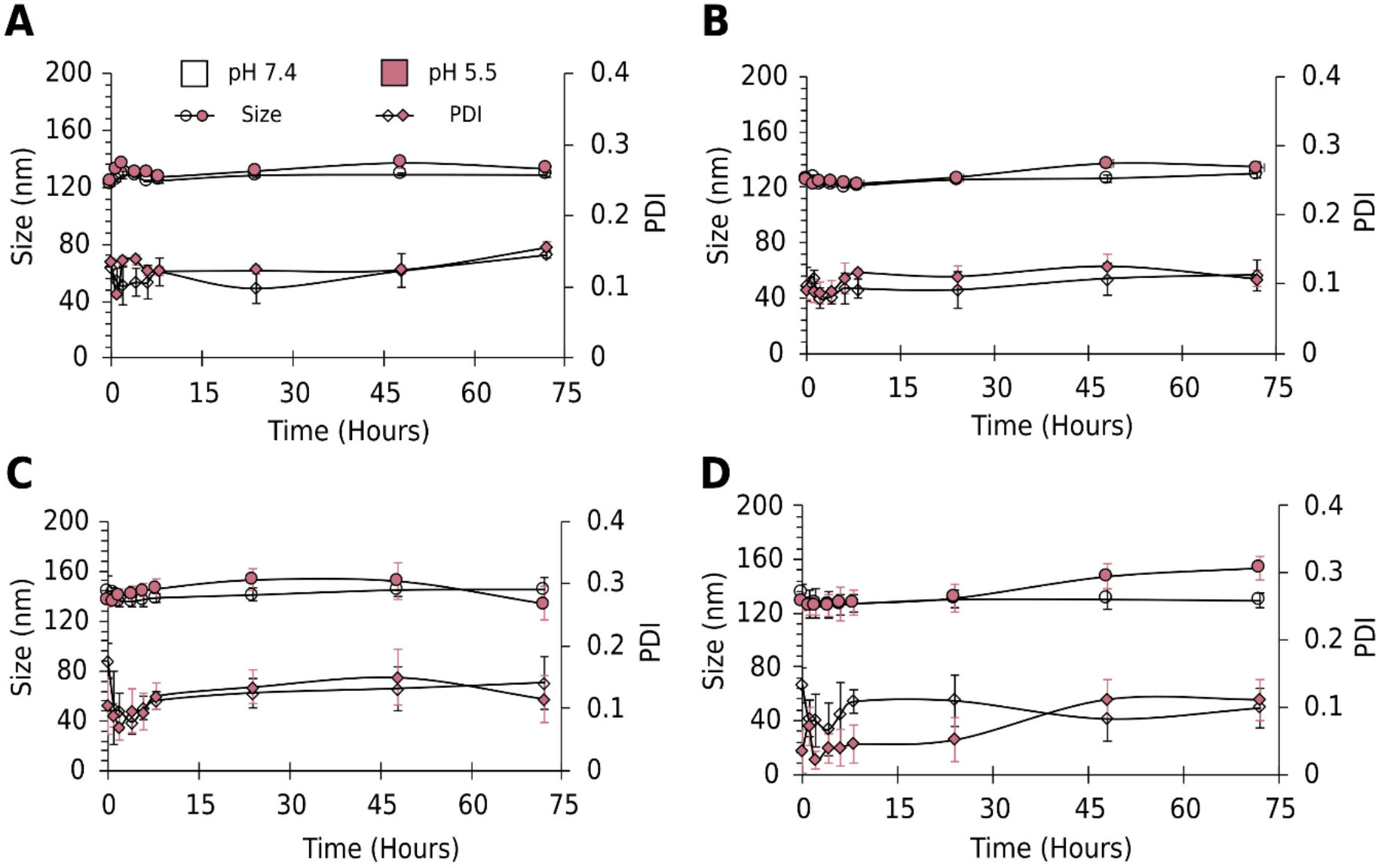
Short-term stability of empty F1 (A), F2 (B), H-F1 (C) and H-F2 (D) in cell culture medium with 10 % FBS at pH 7.4 and pH 5.5. Data were expressed as mean ± standard deviation (SD), n=3. Abbreviations: PDI, polydispersity index.

The size of H-F1 remained stable at pH 7.4, and some fluctuations were observed at pH 5.5 after 24 hours (**Figure 4C**). The reduction in size after interaction with cell culture medium supplemented with serum was observed in H-F2 at 0 hour. Size decreased by approximately 10 nm at pH 7.4 and by approximately 17 nm at pH 5.5 (**Figure 4D**). Additionally, after 24 hours of incubation, the size of H-F2 started to increase till the end of the experiment at pH 5.5. PDI of both hybrid nanoparticles remained below 0.2 at both pH values.

### Encapsulation efficiency and drug release

Following encapsulation, DOX leakage from the liposomes and the hybrid nanoparticles was evaluated at weekly intervals (**Figure 5A**). Generally, F2, containing DPPS, allowed significantly higher DOX encapsulation (60.2 ± 2.3 %), compared to F1 (23.8 ± 0.3 %) (**Figure 5B**). However, hybridization increased the EE% of both formulations, to approximately 45% in F1 and to approximately 76% in F2. At week 4, the amount of DOX retained in F1 liposomes had decreased by nearly 50% during storage, indicating carrier membrane leakiness. In contrast to F1, F2 liposomes showed stable DOX encapsulation for one month without a significant decrease. In H-F1-DOX, the amount of drug retained during storage increased to approximately 70% compared with its non-hybridized counterpart. Similarly to F2-DOX, the amount of DOX encapsulated in H-F2 remained stable over 4 weeks.

**Fig. 5.**
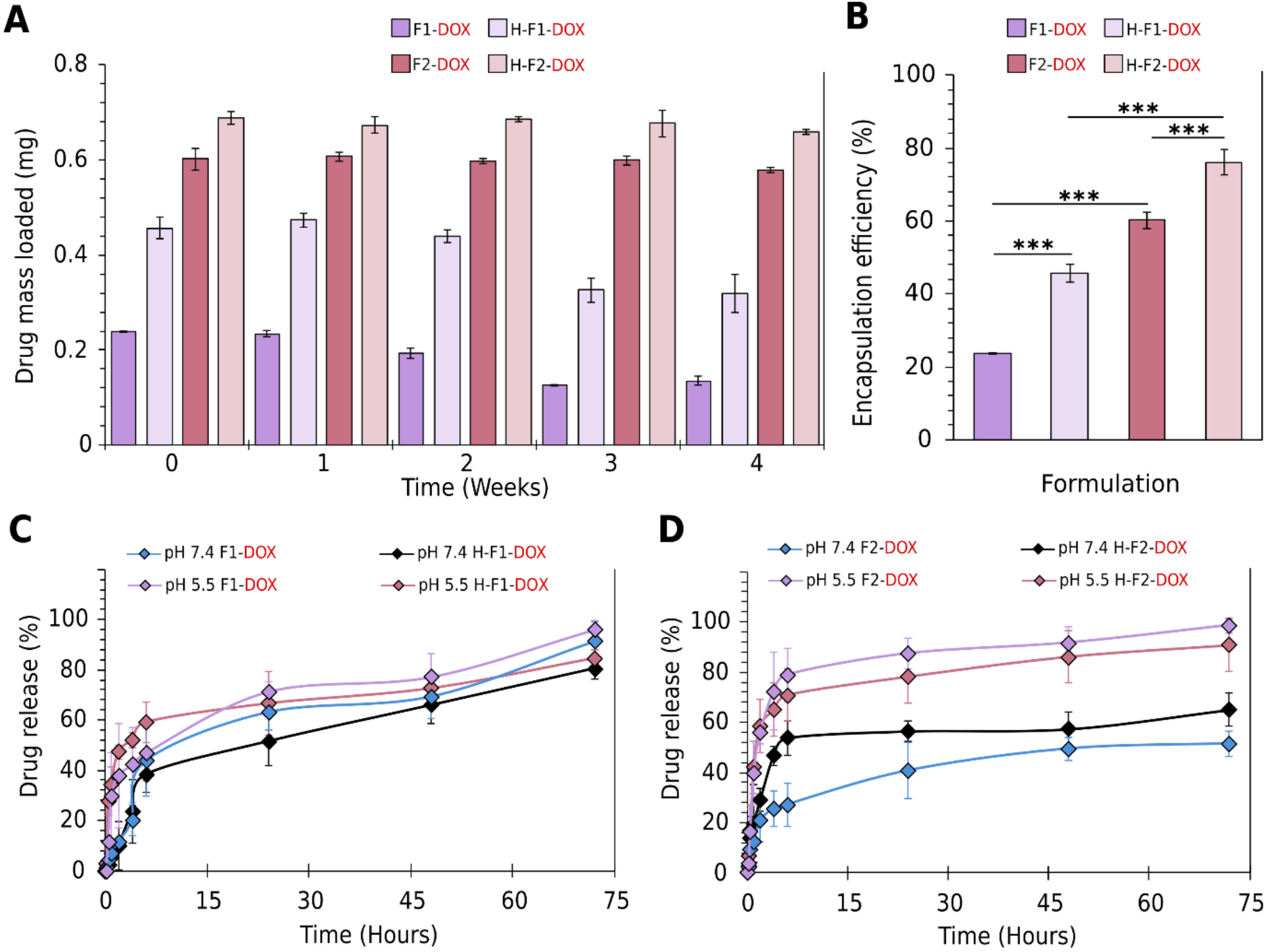
DOX retention (A), Encapsulation efficiency (%) (B) and release profiles of F1 (C) and F2 (D) liposomes and hybrids. Significance level: p < 0.001 (***), one-way ANOVA with Tukey’s post hoc test. Data were expressed as mean ± standard deviation (SD), n=3. Abbreviations: DOX, doxorubicin.

Additionally, cumulative release profiles over 72 hours were quantified in serum-containing cell culture medium at pH 7.4 and pH 5.5. F1 liposomes released encapsulated DOX within 72 hours, similarly under both neutral and acidic conditions (**Figure 5C**). F2 exhibited maximal DOX release at pH 5.5 and almost 50% lower release at pH 7.4. (**Figure 5D**). DOX release profile was influenced by hybridization. In contrast to liposomes, H-F1 and H-F2 showed slower overall DOX release over 72 hours. H-F2-DOX exhibited pH-dependent release profile, but it was decreased in comparison to liposomes.

### Effect of nanoparticles on cell viability

The effects of empty nanoparticles on U-87MG cell viability were evaluated using the MTT assay at 72 h under normoxic and hypoxic conditions (**Figure 6**). To determine whether empty nanoparticles exhibit cytotoxic effects on GBM cells, we tested carrier concentrations ranging from 2.5 to 10 μg/mL. In normoxia, cell viability remained above 85% across all tested groups at the highest nanoparticle concentration (**Figure 6A**). Under hypoxia-mimetic conditions, empty F2 liposomes reduced U-87MG cell viability by an additional 7% compared with normoxia at a concentration of 10 μg/mL. All other empty nanoparticles showed similar effects on cell viability in both normoxic and hypoxic conditions (**Figure 6B**). U-87MG cells’ morphology in carrier treated groups was comparable to untreated controls under both normoxic (**Figure 6C**) and hypoxia-mimetic conditions (**Figure 6D**).

**Fig. 6.**
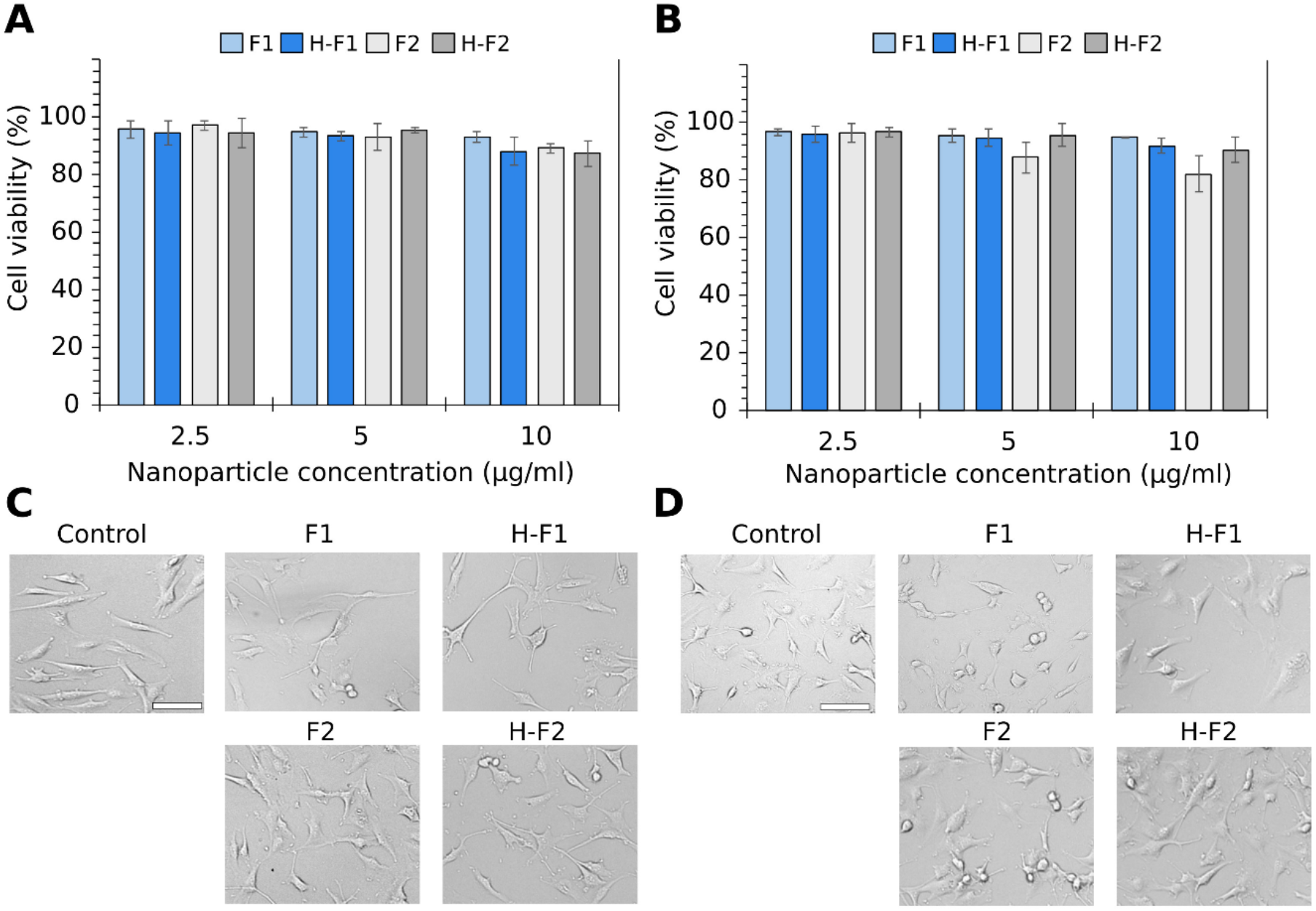
Effect of empty nanoparticles on U-87MG cell viability under normoxia (A,C) and hypoxia-mimetic conditions (B,D). Data were expressed as mean ± standard deviation (SD). Scale bar = 100 μm.

To assess whether DOX encapsulation in the tested nanoparticles influenced its cytotoxic effect, cells were treated with DOX concentrations ranging from 0.00032 to 1 μg/mL. Under normoxic conditions, there were no significant differences between DOX-loaded nanoparticles (F1-DOX, F2-DOX, H-F1-DOX, H-F2-DOX) and free DOX (**Figure 7A**). H-F1-DOX showed the most potent effect on GBM cells, with an EC_50_ value of 0.019 μg/ml (**Figure 7B**). The least potent formulation was H-F2-DOX, with an EC_50_ value of 0.034 μg/ml. However, there were no significant differences between the groups, indicating that the encapsulated nanoparticles exhibited cytotoxic effects comparable to free DOX. Moreover, compared to untreated control cells, treated cells exhibited reduced viability and morphological changes characterized by cell detachment and cell rounding (**Figure 7C**).

**Fig. 7.**
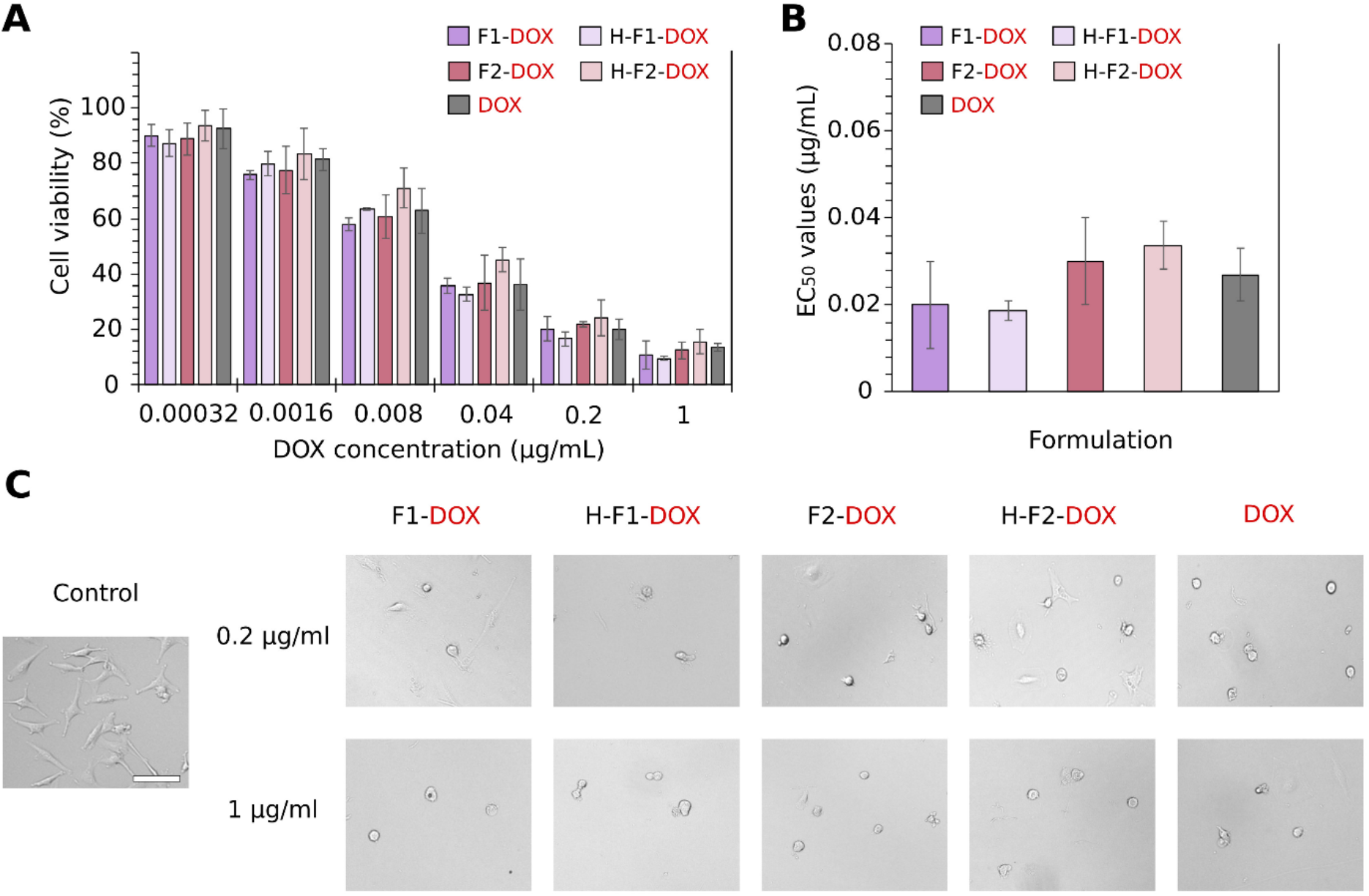
Effect of encapsulated carriers on U-87MG cell viability (A), EC_50_ values of encapsulated carriers and free DOX (B) and morphological changes induced by encapsulated carriers and free DOX (C) under normoxic conditions. One-way ANOVA with Tukey’s post hoc test was used (p < 0.05); no significant differences were observed between groups. Data were expressed as mean ± standard deviation (SD), n=3. Scale bar = 100 μm. Abbreviations: DOX, doxorubicin, EC_50_, half maximal effective concentration.

In contrast to normoxia, under hypoxic conditions, DOX encapsulated into carriers had a significantly higher cytotoxic effect on U-87MG viability, compared to free DOX at concentrations of 1 μg/mL and 0.2 μg/mL (**Figure 8A**). However, there were no significant differences between liposomes and their hybrid counterparts. At 0.04 μg/ml, only F2 and H-F2 had a significant effect on U-87MG cell viability, compared to free DOX. In contrast to normoxia, H-F2-DOX exhibited the lowest EC_50_ value compared to the other tested nanoparticles and free DOX (**Figure 8B**). At a concentration of 0.2 μg/mL, a higher number of attached cells was observed, particularly in the H-F2-DOX group (**Figure 8C**).

**Fig. 8.**
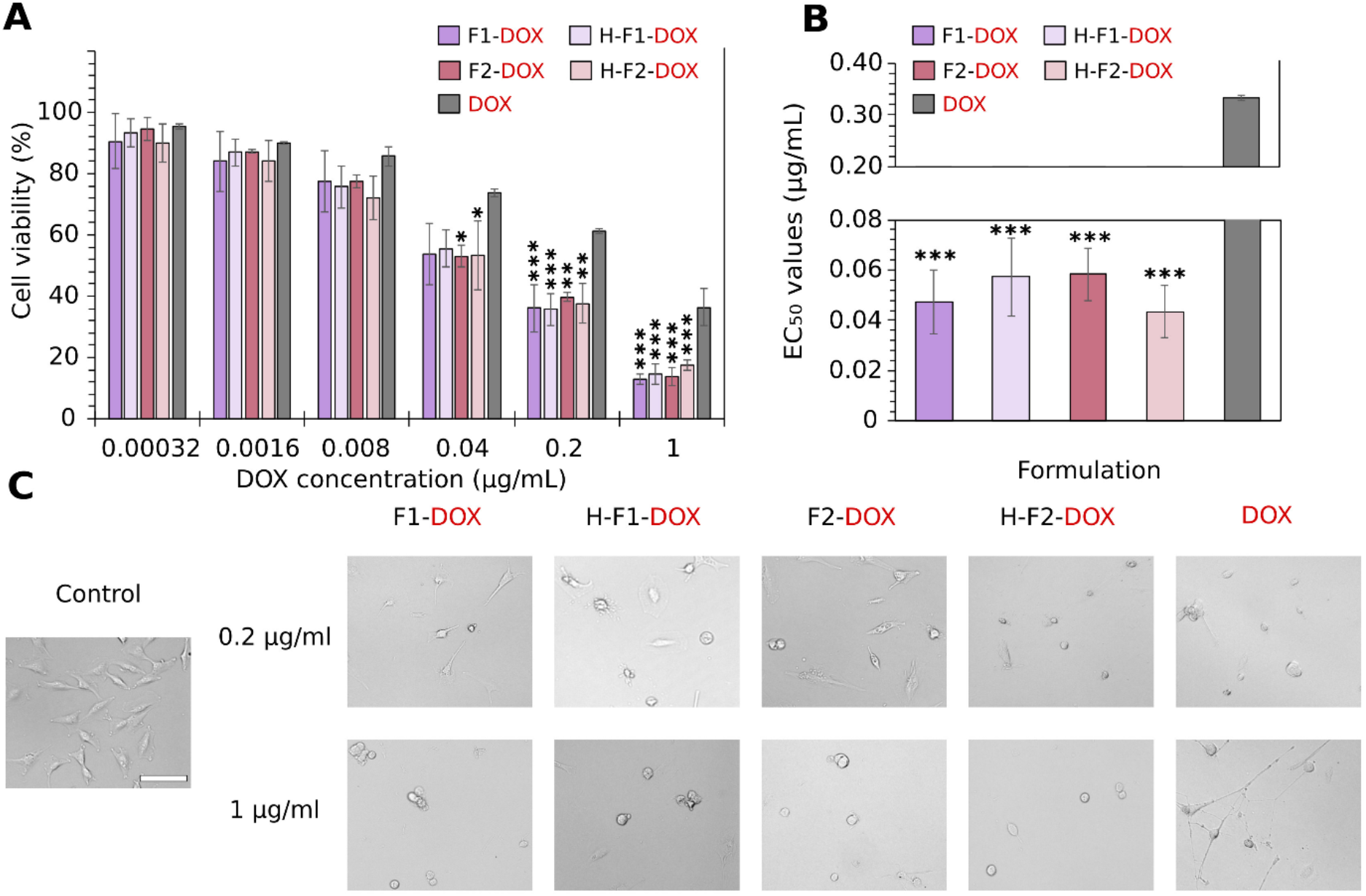
Effect of encapsulated carriers on U-87MG cell viability (A), EC_50_ values of encapsulated carriers and free DOX (B) and morphological changes induced by encapsulated carriers and free DOX (C) under hypoxia-mimetic conditions. Significance levels: p < 0.05 (*), p < 0.01 (**), p ≤ 0.001 (***), one-way ANOVA with Tukey’s post hoc test. Data were expressed as mean ± standard deviation (SD), n=3. Scale bar = 100 μm. Abbreviations: DOX, doxorubicin, EC_50_, half maximal effective concentration.

To further evaluate the data, statistical analysis was performed to compare the cytotoxic effect of each nanoparticle under normoxic and hypoxic conditions (**Figure 9**). Under hypoxic conditions, the highest increase in EC_50_ was observed for H-F1-DOX. No significant differences between normoxic and hypoxic conditions were observed for the remaining nanoparticles. In the case of H-F2-DOX, only a minimal difference was detected (mean difference = −0.0097 ± 0.008), further supporting comparable cytotoxic activity under both conditions.

**Fig. 9.**
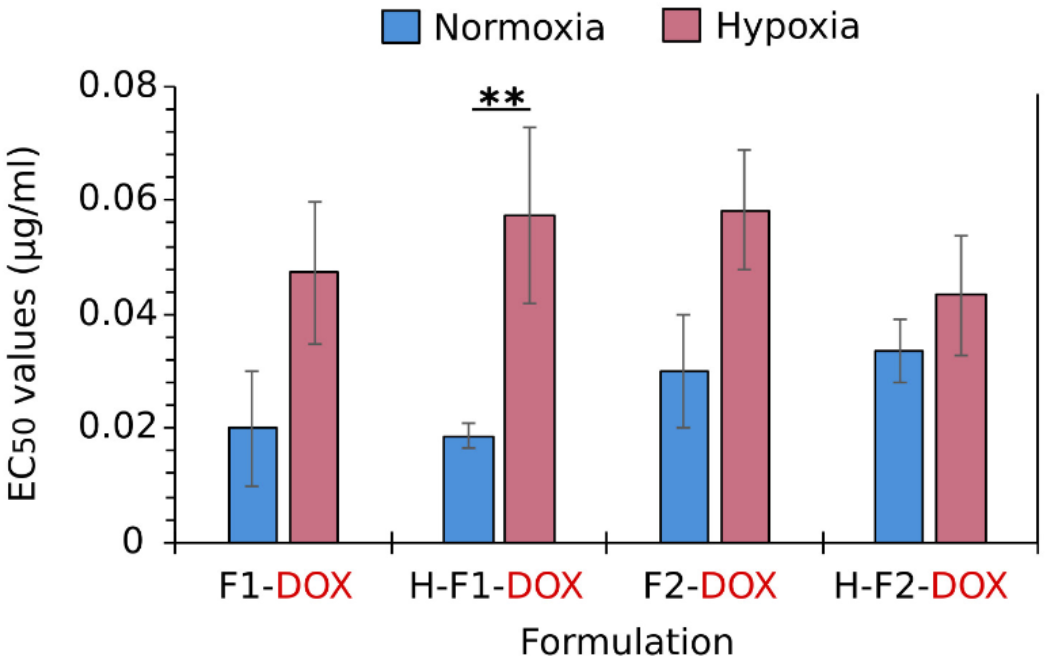
Comparison of nanoparticle cytotoxic effect under normoxic and hypoxic conditions. Significance level: p < 0.01 (**), one-way ANOVA with Tukey’s post hoc test. Data were expressed as mean ± standard deviation (SD), n=3. Abbreviations: DOX, doxorubicin, EC_50_, half maximal effective concentration.

## 4. Discussion

In this study, we synthesized and evaluated two liposomal formulations, DPPC:Chol:DSPEmPEG2000 (F1) and DPPC:DPPS:Chol:DSPEmPEG2000 (F2) and their hybrid derivatives (H-F1 and H-F2) for DOX delivery to GBM cells. In this case, F1 was selected as a less complex formulation that already showed an advantage in the intracellular uptake of gemcitabine and its anticancer activity in Caco-2 cells, compared to other liposomal formulations, such DPPC/Chol/Oleic acid (molar ratio 8:3:1) and DPPC:Chol:DPPS (molar ratio 6:3:1) [48]. F2 additionally includes DPPS, which contains an ionizable carboxylic group (pK_a_^app^≅5.5) that influences stability of the phospholipid at different pH [49]. Both empty and DOX-loaded liposomal formulations were thoroughly characterized alongside EVs-liposome hybrid nanoparticles to establish a controlled baseline for interpreting the effects of membrane hybridization. EVs were collected from J774 cells and purified by size-based chromatographic separation. For this proof-of-concept study, J774 macrophages were selected as a well-characterized cell line with high EVs yield and strong relevance in the literature [50,51]. The average size of selected EVs was within the typical small EVs range [52], with a slightly higher PDI compared to empty liposomes. Previous studies have shown that bioinspired hybrid liposomes can overcome chemoresistance in ovarian cancer [53], exhibit increased accumulation in ischemic brain regions [54] and targeting in breast cancer cells [55], highlighting their versatility arising from the EVs origin. Based on these findings, EVs-liposome hybrid nanoparticles were developed to overcome poor drug accumulation in GBM associated with its aggressive and immune cell–infiltrated microenvironment.

The physicochemical properties of nanoparticles strongly influence the transport cascade involved in targeting GBM [56]. Nanoparticles designed for brain delivery are typically 10–100 nm, however, BBB permeability also depends on properties such as composition and surface charge, with surface functionalization in particular enabling particles larger than 200 nm to cross the BBB [57,58]. Accordingly, unlike bare liposomes, EVs–liposome hybrids are expected to exhibit active targeting properties, and their final size ≤ 200 nm was considered acceptable [59]. Hybrid nanoparticles did not show significant changes in size, indicating that extrusion was successful after freeze-thaw cycling. Empty nanoparticles exhibited a mean size of ~150 nm, with an increase up to 22 nm observed following DOX encapsulation. Such a size change induced by loading was not observed in F1 liposomes, suggesting that EVs and liposomal membrane fusion affects DOX trapping. H-F1-DOX and H-F2-DOX exhibited narrow size distributions and moderately to highly negative zeta potential values, both consistent with reports in the literature on nanoparticles designed for brain delivery [60,61].

Aggregation and flocculation result from colloidal instability and are reflected by changes in particle size, PDI, and zeta potential [62]. Accordingly, these parameters are monitored over time to assess the storage stability of newly developed nanoparticles [63,64,65]. Excellent one-month storage stability was observed for liposomes, independent of DOX loading, as well as for empty nanoparticles. The DOX encapsulation process is sensitive and can destabilize the liposome membrane depending on lipid composition, especially in formulations containing an ammonium sulfate gradient [66]. In our case, hybridization increased membrane sensitivity to drug loading, leading to reduced storage stability of H-F1-DOX and H-F2-DOX. Although a shortened shelf life may limit practical applicability, experimental reproducibility was ensured by using freshly prepared batches [67]. As buffers alone do not fully reflect biological environments, a more complex medium was chosen for short-term stability evaluation [68]. F1 and F2 exhibited an initial decrease in size in serum-supplemented cell medium, regardless of pH. This behavior is consistent with reported serum-induced shrinkage of PEGylated liposomes caused by osmotic pressure driving water out of the liposomal core [69]. In hybrids, this decrease was more pronounced for H-F2 at pH 5.5, consistent with reports showing that acidic conditions enhance protein adsorption [70]. Apart from the initial size decrease, both particle size and PDI of all tested nanoparticles remained stable over 72 hours. These data suggest that H-F1 and H-F2 did not undergo premature breakdown and remained stable under stressful conditions.

Despite comparable behavior of the empty hybrid nanoparticles, the therapeutic hybrid nanoparticles displayed distinct properties. H-F1-DOX and H-F2-DOX exhibited moderate and high EE%, respectively, compared to clinically used liposomal DOX [71]. F1 also showed significantly lower EE% in comparison to F2, suggesting that this formulation was not beneficial for pH-driven DOX trapping. The previously reported gemcitabine loading in F1 is not directly comparable, as the drug association involved substantial bilayer adsorption rather than membrane permeation [48]. The enhanced DOX retention observed in the F2 bilayer was preserved in the H-F2-DOX formulation, suggesting a beneficial role of DPPS in DOX trapping. Notably, hybridization improved DOX EE% and retention in the F1 bilayer, most likely due to EVs components such as cholesterol, sphingomyelin, and phosphatidylserine, which stabilize membrane structure [72,73]. DOX release was evaluated under physiological (pH 7.4) and late endosomal or acidic tumor-associated (pH 5.5) conditions [74,75]. H-F1-DOX and H-F2-DOX displayed a moderate initial release followed by sustained release. Acidic pH enhanced cumulative DOX release for both hybrid nanoparticles, particularly for H-F2-DOX, which may be beneficial for reducing off-target toxicity [75]. Nanoparticle characterization data suggest that hybridization improves DOX trapping, leading to higher drug encapsulation and enhanced retention during storage, however, EVs–liposome hybrids largely preserve the characteristics of their liposomal base, as evidenced by the distinct differences between the two formulations.

After characterization, the initial biological response was assessed in U-87MG cells by MTT assay. As expected, the cell viability assay showed that empty nanoparticles did not exert cytotoxic effects on GBM cells, indicating that the observed anticancer activity can be attributed to the encapsulated cargo [76]. In two-dimensional cultures, many cancer cell lines tend to be more sensitive to treatment due to monolayer growth, making these assays less representative of *in vivo* conditions but well-suited for evaluating the primary effect of nanoparticles on GBM cells [77,78]. Our results showed that the tested nanoparticles had comparable cytotoxic effects to free DOX under normoxic conditions. Unlike free drug molecules, encapsulated drug nanoparticles must first undergo cellular uptake and intracellular release before they can induce cytotoxicity. This mechanism aligns with findings from earlier research on liposomal chemotherapeutics, which often show similar or even lower toxicity compared to free drugs in cancer cell viability tests [79,80,81]. To evaluate whether the nanoparticles remain effective under hypoxia-mimetic conditions, cell viability was assessed in cells treated with CoCl2. Chemically induced hypoxia reduces the sensitivity of cancer cells to chemotherapy by stabilizing HIF-1α. This triggers drug resistance mechanisms, such as increased drug efflux and a metabolic shift toward glycolysis [82]. Both P-gp-mediated efflux and protonation at acidic pH can decrease intracellular DOX levels [83]. In our study, this was reflected by a dramatic increase in the EC_50_ of DOX in U-87MG cells under hypoxia-mimetic conditions compared to normoxia. Previous reports have shown that liposomal DOX does not significantly enhance cytotoxicity compared to free DOX in breast and ovarian cancer cells under hypoxic conditions [84,85]. The beneficial effect in enhancing DOX cytotoxicity on HGC-27 cells in hypoxia was noticed in hypoxia-responsive hybrid liposomes [85]. In our study, all tested nanoparticles showed an advantage in retaining DOX effect on GBM cells in hypoxia-mimetic conditions. In this case, only H-F1-DOX exhibited a significantly weaker cytotoxic effect, while other therapeutic nanoparticles showed comparable results under the tested conditions. It is noteworthy that the tested hybrid nanoparticles behaved differently under CoCl_2_ treatment, as H-F2-DOX exhibited the strongest effect on GBM cells. Given the marked heterogeneity of GBM, including variations in oxygen levels and signalling pathways, it is essential for nanoparticles to maintain consistent anticancer activity across all tumor regions [86]. H-F2-DOX was successfully synthesized, and characterization demonstrated its suitability as nanoparticles for DOX delivery and functionality in GBM cells. A more detailed evaluation of its application in GBM treatment should be performed and will be addressed in future studies.

## 5. Conclusion

In conclusion, this study systematically evaluated the potential of EVs–liposome hybrids for DOX delivery to GBM. Physicochemical characterization demonstrated that H-F2-DOX maintains structural integrity in cell medium with serum at physiological pH and exhibits high EE% and drug retention, supporting its suitability as stable nanoparticles for DOX delivery. Both hybrid nanoparticles exerted strong cytotoxic effects on GBM cell viability under normoxia, and H-F2-DOX demonstrated a clear advantage over H-F1-DOX and free DOX under hypoxia-mimetic conditions. These findings provide a solid foundation for further evaluation of H-F2-DOX in more complex three-dimensional cell culture and BBB models, to evaluate its potential advantages over F2 liposomes in BBB permeation and GBM cell uptake.

## Acknowledgements

The authors thank Simona Tučkutė (Lithuanian Energy Institute, LEI) for valuable assistance with scanning electron microscopy (SEM) analysis of EVs.

## Author Contributions

Conceptualization, V.P. and C.C.; methodology, V.P. and C.C.; validation, V.P.; formal analysis, G.D.; investigation, G.D.; resources, V.P. and C.C.; data curation, G.D.; writing-original draft preparation, G.D.; writing-review and editing, G.D., C.C. and V.P.; visualization, G.D.; supervision, V.P. All authors have read and agreed to the published version of the manuscript.

## Compliance with ethical standards

### Ethical approval

This article does not contain any interventional studies with human participants or animals performed by any of the authors.

